# Single-cell Transcriptomics Reveal Different Maturation Stages and Sublineages Commitment of Human Thymic Invariant Natural Killer T cells

**DOI:** 10.1101/2022.08.10.503443

**Authors:** Kristina Maas-Bauer, Swati Acharya, Jeanette Baker, David B. Lewis, Robert S. Negrin, Federico Simonetta

## Abstract

Invariant natural killer T (iNKT) cells are a subset of heterogenous T-cells with potent cytotoxic and immunomodulatory properties. During thymic development, murine iNKT cells go through different maturation stages and differentiate into distinct sublineages, namely iNKT1, iNKT2, and iNKT17 cells. To define maturation stages and to assess sublineage commitment of human iNKT cells during thymic development, we performed single-cell RNA sequencing analysis on human thymic iNKT cells. We show that these iNKT cells displayed heterogeneity and unsupervised analysis identified two clusters: one with an immature profile with high expression of genes that are important for iNKT cell development and enriched in cells expressing an iNKT2 signature, whereas a second cluster displayed a mature, terminally differentiated profile resembling murine iNKT1 cells. Trajectory analysis suggested an ontological relationship between the two clusters. Our work provides the first single cell transcriptomic analysis of thymic human iNKT cells offering new insights into their developmental process in humans.

## Introduction

Invariant Natural Killer T (iNKT) cells are a rare subset of innate lymphocytes representing less then 1% of the total lymphocyte population both in humans and mice. iNKT cells express a semi-invariant TCR, consisting of a Vα14Jα18 chain paired with a limited selection of beta chains in mice and Vα24Jα18 typically pairing with Vβ11 in humans, that recognize glycolipids presented in the context of the non-polymorphic, MHC-like molecule CD1d (Kawano et al, 1997; Lantz & Bendelac, 1994). Upon stimulation, iNKT cells can promptly release a wide range of cytokines, allowing to iNKT cells to exert a spectrum of pleiotropic functions, ranging from antitumor effects to immune-regulatory activity (Crosby & Kronenberg, 2018; Matsuda et al, 2008).

Traditionally, thymic development of murine iNKT cells has been defined by different maturation stages based on the surface expression of CD24, CD44 and NK1.1: CD24^hi^CD44^lo^, NK1.1^lo^ immature precursor iNKT cells (iNKT0 or iNKTp; (Benlagha et al, 2002); CD24^lo^CD44^lo^, NK1.1^lo^ stage 1 iNKT cells; CD24^lo^CD44^hi^, NK1.1^lo^ stage 2 iNKT cells; CD24^lo^CD44^hi^, NK1.1^hi^ stage 3 iNKT cells. Interestingly, these maturation stages have been associated with different functional properties with stage 1 and 2 iNKT cells producing interleukin 4 (IL-4) and interleukin10 (IL-10), stage 3 iNKT cells producing interferon-γ (IFN-γ) and having limited proliferation potential compared to stage 1 and 2 iNKT cells (Coquet et al, 2008; Gadue & Stein, 2002; Pellicci et al, 2002). Such model has been challenged by recent reports based on single cell genomic analyses identifying the coexistence of different maturation iNKT stages in the murine thymus and also suggesting that populations with mixed characteristics exist (Harsha Krovi et al, 2020).

Partly opposed to the theory of different maturation stages, several studies indicated the existence of at least three, terminally differentiated, murine iNKT sublineages, namely Th-1 like iNKT (iNKT1), Th-2 like iNKT (iNKT2) and Th-17 like iNKT (iNKT17) cells (Engel et al, 2016; Lee et al, 2013). These subsets are characterized by the differential expression of the transcription factors promyelocytic leukaemia zinc finger (PLZF), GATA Binding Protein 3 (GATA3), T-bet and RAR-related orphan receptor gamma (RORγt) (iNKT1: PLZF^lo^, T-bet^+^, iNKT2: PLZF^hi^ GATA3^hi^, iNKT17: PLZF^int^RORγt^+^, and PLZF^lo^, T-bet^+^) (Engel et al, 2016; Lee et al, 2013), produce a unique cytokine profile (Lee et al, 2013) and have special distribution in tissues (Lee et al, 2015). We recently demonstrated that distinct iNKT sublineages exert different functions, iNKT2 and iNKT17 displaying immunoregulatory properties while iNKT1 exerting the strongest cytotoxic activity (Maas-Bauer et al, 2021). Importantly, in contrast to what observed in conventional T cells whose Th1, Th2 or Th17 lineage commitment takes place upon antigen encounter in the periphery, iNKT cell sublineages differentiation already takes place at the thymic level (Engel et al, 2016; Lee et al, 2013). Importantly, a recent single-cell report on thymic iNKT cells of pre-pubertal pigs showed big differences between murine and porcine iNKT cells (Gu et al, 2022). In this study porcine iNKT cells were unexpectedly homogeneous, with 97% of cells expressing an iNKT2 genotype.

Despite the progresses in the understanding of animalistic iNKT cell development and differentiation, little is still known about human iNKT cell thymic development. Studies investigating maturation processes of human iNKT cells based on phenotypic analysis have shown that, similarly to murine iNKT cells, the predominant iNKT cell population in neonatal human thymus are CD4^+^CD161^-^ iNKT cells, whereas CD4^-^CD161^+^ iNKT cells accumulate with age (Berzins et al, 2005; Sandberg et al, 2004). Whether these subsets of human thymic iNKT cells correspond to distinct maturation stages is still unclear. Moreover, it is still unknown whether human iNKT cells commit at the thymic level toward any of the sublineages profiles reported in mice.

## Results

### Single-cell RNA sequencing identifies two different iNKT cell clusters corresponding to distinct developmental stages in the human thymus

We performed single cell RNA-sequencing (scRNA-seq) on human thymic iNKT cells FACS-sorted from thymi from 3 newborn/infant donors (Figure 1A). After removing doublets (>2500 gene counts), cells with lowest (<200) gene counts and cells with high (>5%) mitochondrial gene content, 341 cells were retained for downstream analysis. Unsupervised Uniform Manifold Approximation and Projection (UMAP) analysis segregated human thymic iNKT cells into 2 clearly defined clusters: cluster 0 and cluster 1 (Figure1B). Such distribution was conserved across the three samples originating from the three different donors (Supplemental Figure 1). Differential expression analysis revealed 259 differentially expressed genes between the two clusters. Among the 20 most differential expressed genes (Figure 1C), iNKT cells in cluster 0 were enriched for genes previously reported in mice to be associated with development of iNKT cell precursors, including SRY-Box Transcription Factor 4 (SOX4; (Malhotra et al, 2018)), lymphoid enhancing-factor 1(LEF1; (Berga-Bolaños et al, 2015; Carr et al, 2015)), special AT-rich sequence-binding protein-1(SATB1; (Kakugawa et al, 2017)), and Integral Membrane Protein 2A (ITM2A; (Baranek et al, 2020; Harsha Krovi et al, 2020)). In particular SOX4, encoding the transcription factor important for the development of iNKT cells (Malhotra et al, 2018), was preferentially expressed in cluster 0 and was barely detectable in cluster 1 (Figure 1D). Conversely, cells in cluster 1 expressed genes associated with iNKT full differentiation (Killer Cell Lectin Like Receptor B1 (KLRB1), encoding the surface molecule CD161; Figure 1E). Based on these data we hypothesized that the two clusters could correspond to two different stages of human iNKT development. To address this hypothesis, we performed a pseudotime analysis with cluster 0 defined as the root. This analysis revealed a cell transition trajectory from cluster 0 to cluster 1 (Figure 2A). Along such trajectory we observed a progressive downregulation of the genes encoding the transcription factors LEF1, SATB1 and SOX4 (Figure 1B). This was mirrored by the upregulation of genes known to be involved in iNKT terminal differentiation (KLRB1, (Baranek et al, 2020; Pellicci et al, 2002) and production of cytotoxic molecules (GZMA). Collectively, these data demonstrate the heterogeneity of human thymic iNKT cells and point to the existance of at least two different iNKT maturation stages at the thymic level.

**FIGURE 1.**
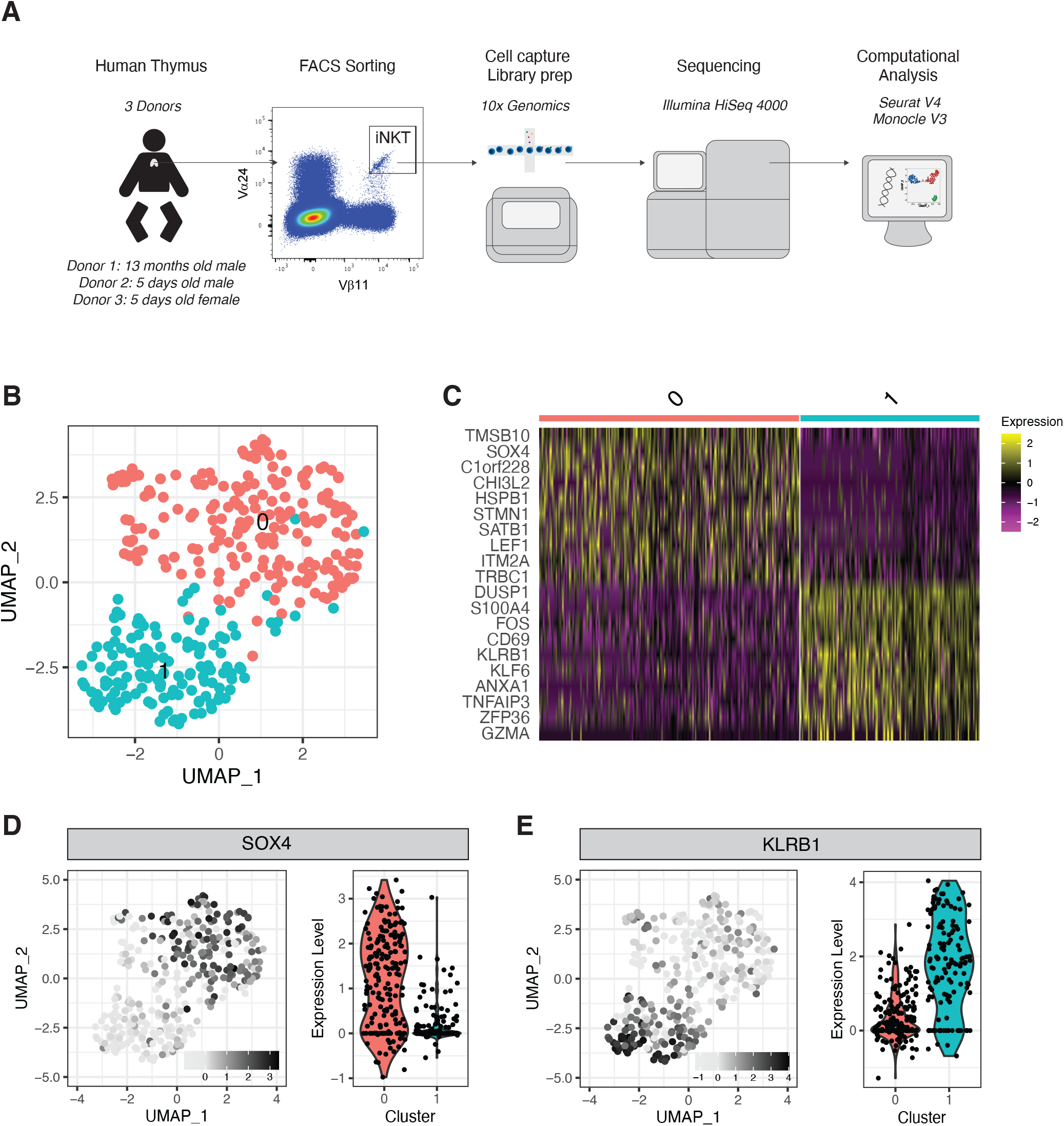
Single-cell transcriptomic analysis reveals heterogeneity of human thymic iNKT cells. **(A)** Schematic representation of the experimental pipeline. **(B)** Uniform Manifold Approximation and Projection (UMAP) plot of scRNA-seq data showing distinct clusters of human thymic iNKT cells. **(C)** SingleLJcell heatmap representing the 10 most highly differentially expressed genes in human thymic iNKT cell clusters. Expression for each gene is scaled (zLJscored) across single cells. **(D-E)** Relative expression and normalized counts of SOX4 **(D)** and KLRB1 **(E)** in human thymic iNKT cell clusters.

**FIGURE 2.**
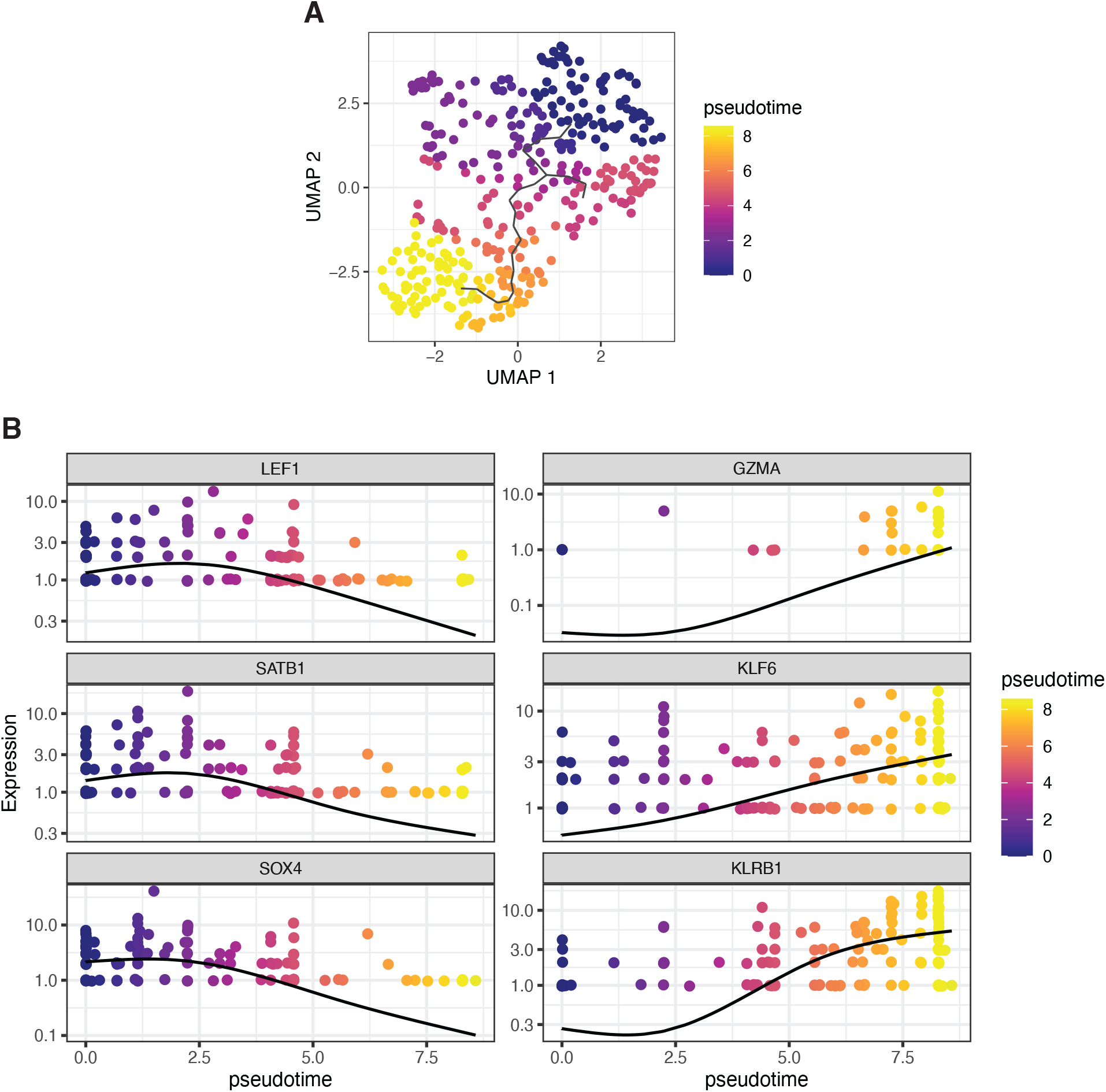
Pseudotime analysis indicates ontological relationship between human thymic iNKT cell clusters. **(A)** UMAP plot of scRNA-seq data colored by pseudotime. **(B)** Single genes normalized expression levels across the pseudotime trajectory. Cells are colored by pseudotime.

### Human iNKT cells with iNKT1 and iNKT2 gene signature are present at the thymic level

Murine iNKT cells have been shown to differentiate at the thymic level into three sublineages, namely iNKT1, iNKT2, and iNKT17 cells, displaying different transcriptomic (Baranek et al, 2020; Engel et al, 2016), epigenomic (Engel et al, 2016) and functional (Maas-Bauer et al, 2021) characteristics. We therefore investigated whether human thymic iNKT cells also displayed signs of sublineage commitment at the thymic level. To this aim, we generated subset-specific gene signatures for iNKT1, iNKT2 and iNKT17 based on human gene orthologues of genes we previously identified in single cell transcriptomic analysis of murine thymic iNKT cells (Maas-Bauer et al, 2021). The composite expression score of iNKT sublineages specific gene sets was calculated using the Seurat’s AddModuleScore function. Our analysis detected an enrichment in iNKT1-signature expressing cells in cluster 1 whereas these cells were barely detectable cluster 0 cells (Figure 3, upper panels). Conversely, cells in cluster 0 displayed an enrichment for iNKT2-signature expressing cells, although such signature was also detectable in some cluster 1 cells (Figure 3, middle panel). We did not identify any clear enrichment of iNKT17-signature expressing cells in either cluster 0 or cluster 1 (Figure 3, bottom panel). To further validate the iNKT-sublineages signatures we generated, we next assessed them on publicly available scRNA-seq data of iNKT cells isolated from human peripheral blood (Erkers et al, 2020). Our analysis clearly identified a cluster of iNKT cells displaying an enrichment in iNKT1-signature genes (Supplemental Figure 2A,B) while we observed the presence of some cells enriched in iNKT2- and iNKT17-signature genes not associated with any specific cluster. In conclusion, our analysis revealed the existence of a population of human iNKT cells committed toward an iNKT1-like, and at a lesser extent iNKT2-like, differentiation during thymic development.

**FIGURE 3.**
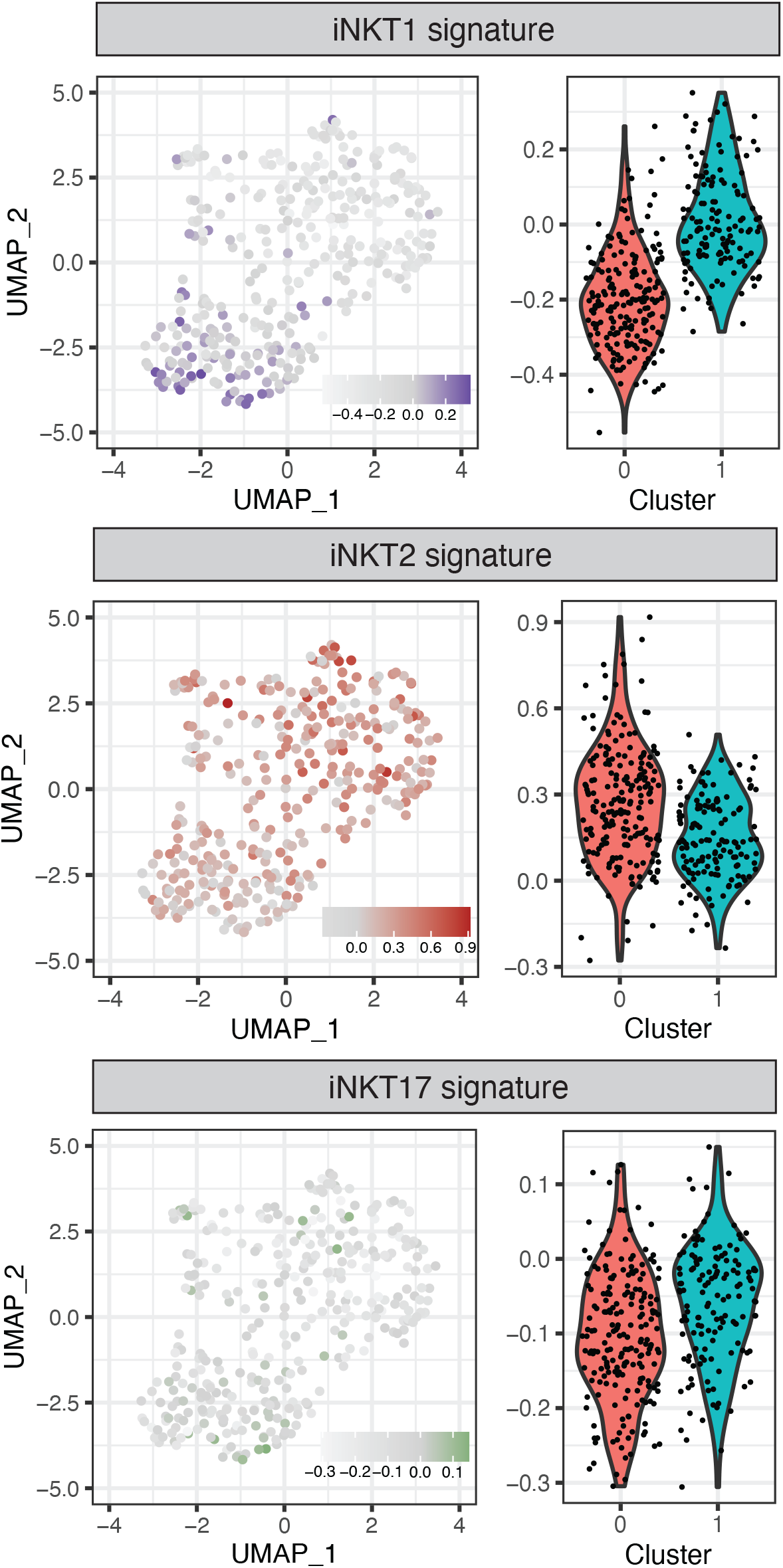
Identification of human thymic iNKT cells expressing an iNKT1 and iNKT2 signature. Relative expression and normalized expression of iNKT1 (upper panels), iNKT2 (middle panels) and iNKT17 (lower panels) signatures. Color intensity represents the composite expression score of iNKT sublineage-specific gene sets as calculated using the Seurat’s AddModuleScore function.

## Discussion

In this study, we used single-cell transcriptomics to analyze human thymic iNKT cells maturation processes and to investigate if, similarly to their murine counterpart, human iNKT cells undergo sublineage differentiation at the thymic level. Our analysis revealed the trajectory of human iNKT cells development through two different maturation stages. More importantly, we provide for the first time evidence that human iNKT cells commit to an iNKT1-sublineage differentiation already at the thymic level.

Unsupervised clustering revealed two clusters of human thymic iNKT cells that mainly differed for the expression of genes encoding for proteins involved in iNKT development and differentiation. The RNA for the transcription factors SOX4 and LEF1 and for the proteins ITM2A and SATB1 was more highly expressed in Cluster 0. SOX4 and LEF1 have been shown to be important transcription factors for iNKT cell development (Malhotra et al, 2018). LEF1, for example, is a crucial transcription factor for iNKT cell proliferation (Berga-Bolaños et al, 2015; Carr et al, 2015)and two extensive studies of murine thymic iNKT cells showed lately that LEF1 and SOX4 are highly upregulated in NKT0 cells (Baranek et al, 2020; Harsha Krovi et al, 2020). In line with our findings, ITM2a was also highly upregulated in the NKT0 population in these analyses (Baranek et al, 2020; Harsha Krovi et al, 2020). ITM2A is a target gene of GATA-3, a transcription factor involved in differentiation of murine iNKT cells and especially for the production of T_h_2 cytokines by iNKT cells (Kim et al, 2006). Moreover, SATB1, a protein involved in the regulation of iNKT cell development in mice (Kakugawa et al, 2017), was also upregulated in cluster 0 cells. Regarding cluster 1, we found that genes encoding for cytotoxic molecules like GZMA (encoding for granzyme A) were enriched. Moreover, the gene encoding KLRB1, a molecule associated with effector function in several lymphocytic subset (Takahashi et al, 2006) was also expressed at higher levels in cluster 1 cells compared to cells in cluster 0. Cluster 1 cells expressed higher levels of CD69 transcript, encoding a surface molecule associated with activated state in T cells (Sancho et al, 2005) and augmented cytotoxicity in NK cells (Clausen et al, 2003; Moretta et al, 1991). Overall, the gene program of cells in cluster 0 is reminiscent of murine iNKT0/p cells with upregulation of genes involved in iNKT cell development and proliferation, whereas cells in cluster 1 resembled murine mature iNKT cells with an upregulation of molecules responsible for effector function and associated with an activated phenotype. Trajectory analysis with pseudo-time of the same population (Figure 2), revealed a dynamic process of cells in cluster 0 maturing to cells in cluster 1(Figure 2B), resulting in a cytotoxic phenotype of the differentiated cells. Taken these findings together, we suggest that different maturation stages of iNKT cells exist in human thymus. These findings are in line with previously reported murine data showing that different maturation stages of iNKT cells are present at the thymic level (Baranek et al, 2020; Harsha Krovi et al, 2020).

We previously showed that murine thymic iNKT1, iNKT2 and iNKT17 cells not only differ in their transcriptomic and epigenetic profile but also exert different functions in vitro and in vivo (Maas-Bauer et al, 2021). To investigate whether human iNKT cells also possess properties attributed to murine iNKT1, iNKT2 and iNKT17 cells, we evaluated the enrichment for human gene orthologues to the ones we previously reported for identification of iNKT-sublineages in mice (Maas-Bauer et al, 2021). We found that cells in cluster 1 are enriched for cells that resemble murine iNKT1 cells, whereas cells in cluster 0 are enriched for cells with an iNKT2 signature, although, at a lesser extent, iNKT2-like cells were also found in cluster 1. This heterogeneity are in line with recent data suggesting that iNKT2 cells are not terminally differentiated and consist of several subpopulations that differ in their maturation grade. During maturation some iNKT2 subsets (iNKT2b) acquire markers usually attributed to iNKT1 cells (Baranek et al, 2020; Harsha Krovi et al, 2020).

Interestingly, iNKT cells with an iNKT17 profile did not cluster in one specific region in the UMAP analysis (Figure 3). Thus, we cannot conclude that iNKT17 cells also exist in human thymus. As human iNKT cells that express RORγt and IL-17 have been described in peripheral blood (Venken et al, 2019), it is possible that they either differentiate in the periphery (Wang & Hogquist, 2018) or emigrate directly after differentiation. In addition to this data, several studies suggest that CD69 expression of iNKT1 cells is responsible for thymic retention (Baranek et al, 2020; Nakayama et al, 2002) and that only iNKT2, iNKT17 and some iNKT1 subsets leave the thymus (Baranek et al, 2020). The iNKT1-like cells in our analysis expressed CD69, which might explain the relatively big proportion of these cells in human thymus.

Before iNKT subsets in mice were investigated, data suggested that iNKT cells are defined by their maturation stage (Pellicci et al, 2002; Watarai et al, 2012). Later iNKT1, iNKT2 and iNKT17 cells were identified as fully differentiated cell populations (Hogquist & Georgiev, 2020; Lee et al, 2013)). More recent reports performed scRNA-seq and showed with trajectory analysis that iNKT cells in the thymus differ in their maturation stage (Baranek et al, 2020; Harsha Krovi et al, 2020). Moreover, they showed that iNKT1 and iNKT2 cells consist of different subgroups and suggest a plasticity at least in iNKT0/p and in the more immature subgroups of iNKT2 cells (Baranek et al, 2020; Harsha Krovi et al, 2020). Thus, owing to these studies it is now possible to reconcile the discovery of iNKT1, iNKT2 and iNKT17 subsets with the knowledge of different maturation stages of iNKT cells in the thymus, suggesting that iNKT2 cells display immature characteristics compared to iNKT1 and iNKT17 cells and probably function as a precursor for these subsets (Baranek et al, 2020; Harsha Krovi et al, 2020). Moreover, a recent study of thymic porcine iNKT cells also demonstrated, that some iNKT2 cell subsets of pre-pubertal pigs had immature properties with an upregulation of LEF1 and SATB1 that are typical for immature iNKT cells (Gu et al, 2022). Intriguingly, these data are also in line with our findings that cells with an iNKT2 signature are present in the human thymus, yet appear with immature properties that also resemble murine iNKT0/p cells. Importantly, these findings are also in line with a study of thymic porcine iNKT cells demonstrating, that some iNKT2 of pre-pubertal pigs had immature properties and upregulated LEF1 and SATB1 that are typical for immature iNKT cells (Quelle).

As the composition of iNKT cells is altered with aging (Papadogianni et al, 2020) and might differ among individuals, our analysis represents a snapshot of iNKT cell distribution at very young age. Furthermore, the number of cells in our analysis is limited due to the paucity of iNKT cells in human thymus and the difficulties to access vital thymic tissue of older individuals. However, to our knowledge this is the first study investigating the transcriptomic differences of human thymic iNKT cells, with respect to iNKT maturation stages and cell subsets. Moreover, our data are in line with recent murine studies showing that iNKT cells with iNKT2 properties have immature characteristics and can give rise to terminally differentiated iNKT1 cells. Our work is a first step to a better understanding of human iNKT cell heterogeneity at the thymic level. Future studies will be needed to explore human iNKT subsets at different ages as well as in peripheral organs to decipher their function.

## Acknowledgments

The authors thank Dr. Dhananjay Wagh at the Stanford Functional Genomics Facility for his excellent technical assistance in the genomics analysis

## Author Contributions

F.S., K.M.-B., and R.S.N. conceived and designed the study; K.M.-B., S.A., F.S., performed the experiments; F.S. analyzed the data; F.S., K.M.-B., and R.S.N. contributed to the interpretation of results; K.M.-B., S.A., F.S., and R.S.N. wrote the manuscript; S.A, J.B and D.B.L provided essential reagents and methods; F.S. and R.S.N. supervised the research. All authors read and approved the submitted version of the manuscript.

## Star Methods

### Resource Availability

#### Lead Contact

Further information and requests for resources and reagents should be directed to and will be fulfilled by the lead contact, Federico Simonetta.

#### Materials availability

This study did not generate new unique reagents.

#### Data and code availability

Single-cell RNA-seq data have been deposited at GEO and will be publicly available as of the date of publication. Accession numbers are listed in the key resources table. DOIs are listed in the key resources table. Any additional information required to reanalyze the data reported in this paper is available from the lead contact upon request.

### Experimental model and subject details

#### Human samples

Thymi were obtained after being removed from 3 newborn/infant donors undergoing cardiac surgery between 5 days and 13 months of age for cardiac diseases without evidence of immunological diseases. The analysis of human thymus samples obtained as surgical waste was reviewed and approved by Stanford University, Institutional Review Board (IRB Protocol #16877; directors Drs. David B. Lewis and Swati Acharya, Pediatric Cardiovascular Surgery, Stanford Children’s Health). Single cell suspensions were obtained by mechanical dissociation, washed, cryopreserved in fetal calf serum and 10% DMSO and stored in liquid nitrogen until use.

### Method Details

#### Preparation of thymic iNKT cells

Thymocytes were thawed at room temperature in RPMI 1640 containing FCS 30%, Penicillin-Streptomycin 1%, DNase I (10 ug/ml, Sigma), and heparin 20 U/ml. After washing, FACS surface staining was performed on ice, after FC blockage, using the following antibodies: Vα24 (FITC, Beckman Coulter), Vβ11 (APC, Beckman Coulter), CD3 (BV605, Biolegend), CD14 (APC Cy7, Biolegend), CD19 (APC Cy7, Biolegend), LIVE/DEAD™ Fixable Aqua Dead Cell Stain Kit (Invitrogen). CD3+, Vα24+, Vβ11+, CD14-, CD19-live cells were FACS-sorted on a FACS Aria-III (BD Biosciences).

#### Single cell RNA sequencing

Single-cell libraries were generated from FACS-sorted human thymic iNKT cells using the Chromium Controller Single-Cell Instrument and Chromium Single Cell 3′ Library & Gel Bead Kit v2 (10x Genomics). Sample demultiplexing, barcode processing, alignment to the GRCh38 assembly of the human genome, and single-cell 3′ gene counting were performed using the Cell Ranger Suite version 2.1. The gene-barcode matrix contained 397 cells, with 383784 mean reads per cell and 1099 (984-1203) median genes per cell. Cells with unique gene counts <200 or >2500, as well as cells with >5% of mitochondrial genes were excluded from the analysis. scRNA-seq analysis was performed using the Seurat R package, version 4. Pseudotime analysis was performed using the Monocle3 package. Previously published scRNA-seq datasets from human iNKT cells recovered from the peripheral blood (PRJNA563899, PRJNA565590; (Erkers et al, 2020)) were analyzed using the same pipeline.

### Key resources table

| REAGENT or RESOURCE | SOURCE | IDENTIFIER |
| --- | --- | --- |
| **Antibodies** |  |  |
| FITC-labeled TCR Vα24 | Beckman Coulter | IM1589 |
| APC-labeled TCR Vβ11 | Beckman Coulter | A66905 |
| BV605-labeled anti-human CD3 Antibody | Biolegend | 31732 |
| APC Cy7-labeled anti-human CD14 Antibody | Biolegend | 398707 |
| APC Cy7-labeled anti-human CD19 Antibody | Biolegend | 302217 |
| LIVE/DEAD™ Fixable Aqua Dead Cell Stain Kit | Invitrogen | L34957 |
| **Biological samples** |  |  |
| Human thymic sampes | Pediatric Cardiovascular Surgery | n/a |
| **Critical commercial assays** |  |  |
| Chromium Single Cell 3' Library & Gel Bead Kit v2 | 10x Genomics | PN-120237 |
| Chromium Controller Single-Cell Instrument | 10x Genomics |  |
| **Deposited data** |  |  |
| **Raw data** | **This paper** | Single-cell RNA-seq data have been deposited at GEO and will be publicly available as of the date of publication. |
| Human iNKT cells in peripheral blood | (Erkers et al, 2020) | NCBI Sequence Read Archive: PRJNA563899, PRJNA565590 |
| Software and algorithms |  |  |
| **Cell Ranger Suite version 2.1** | **10x Genomics** |  |
| **Seurat R package, version 4** | **Stuart et al, 2019** | https://satijalab.org/seurat/ |
| Monocle3 | Trapnell et al., Nature Biotechnology, 2014 | https://cole-trapnell-lab.github.io/monocle3/ |

## Supplemental figure legends

**SUPPLEMENTAL FIGURE 1.**
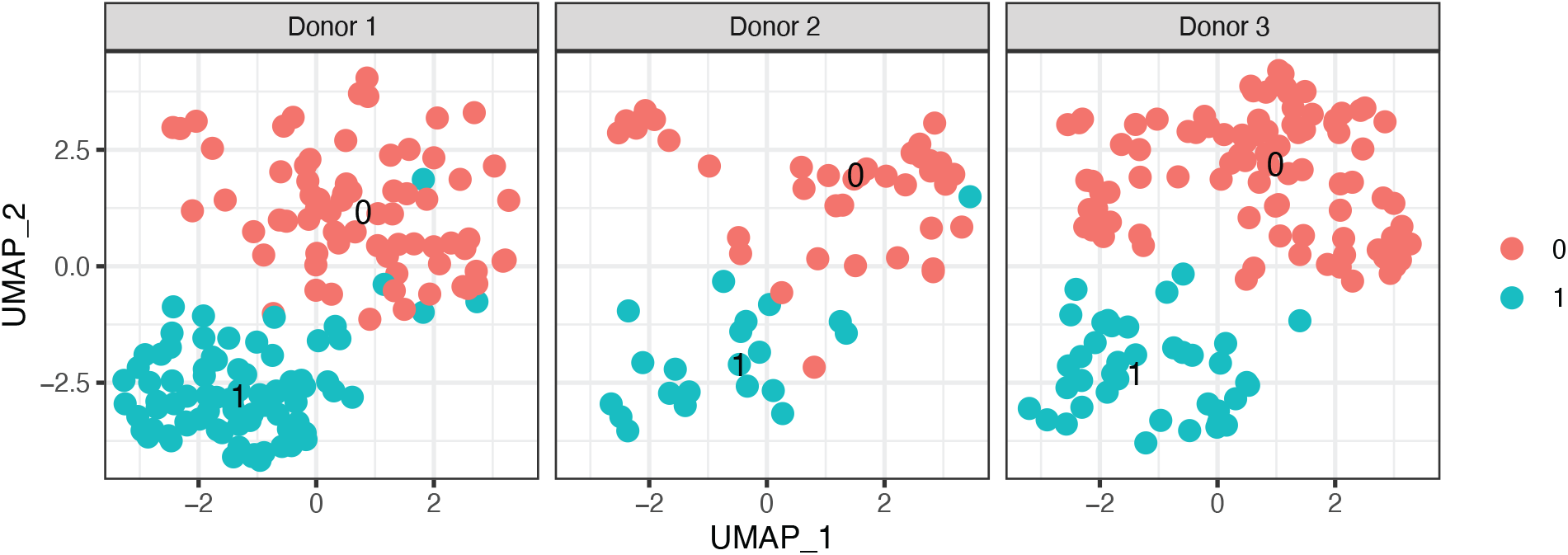
Reproducibility of single-cell transcriptomic analysis of human thymic iNKT across different donors. UMAP plots of scRNA-seq analysis of human thymic iNKT cells splitted by different donor source.

**SUPPLEMENTAL FIGURE 2.**
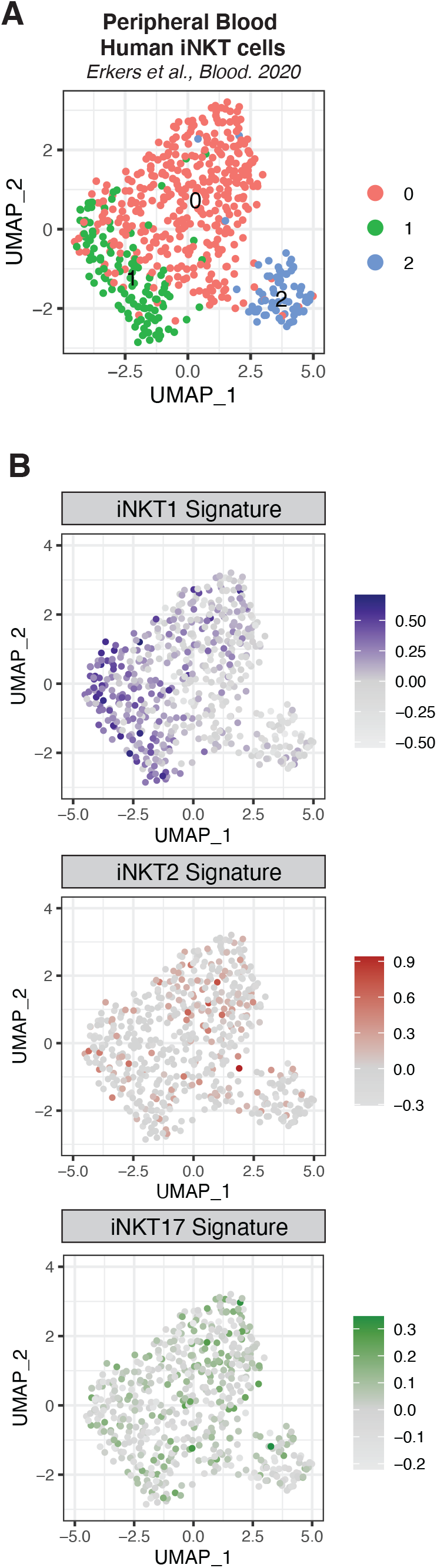
iNKT-sublineage specific signatures expression in peripheral blood human iNKT cells. **(A)** UMAP) plot of publicly available scRNA-seq data (Enkers et al., Blood 2020) showing distinct clusters of human iNKT cells from peripheral blood. **(B)** Relative expression and normalized expression of iNKT1 (upper panels), iNKT2 (middle panels) and iNKT17 (lower panels) signatures. Color intensity represents the composite expression score of iNKT sublineage-specific gene sets as calculated using the Seurat’s AddModuleScore function.

